# A quarter-million-year-old polymorphism drives reproductive mode variation in the pea aphid

**DOI:** 10.1101/2022.10.24.513443

**Authors:** M. Rimbault, F. Legeai, J. Peccoud, L. Mieuzet, E. Call, P. Nouhaud, H. Defendini, F. Mahéo, W. Marande, N. Théron, D. Tagu, G. Le Trionnaire, J.-C. Simon, J. Jaquiéry

## Abstract

Although asexual linages evolved from sexual lineages in many different taxa, the genetics of sex loss remains poorly understood. We addressed this issue in the pea aphid *Acyrthosiphon pisum,* whose natural populations encompass lineages performing cyclical parthenogenesis (CP) and producing one sexual generation per year, as well as obligate parthenogenetic (OP) lineages that can no longer produce sexual females but can still produce males. A SNP-based, whole-genome scan of CP and OP populations sequenced in pools (103 individuals from six populations) showed that a single X-linked region controls the variation in reproductive mode. This 840-kb region is highly divergent between CP and OP populations (F_ST_ = 34.9%), with >2000 SNPs or short Indels showing a high degree of association with the phenotypic trait. Comparison of *de novo* genome assemblies built from long reads did not reveal large structural rearrangements between CP and OP lineages within the candidate region. This reproductive polymorphism still appears relatively ancient, as we estimated its age at ~0.25 million years from the divergence between *cp* and *op* alleles. The low genetic differentiation between CP and OP populations at the rest of the genome (F_ST_ = 2.4%) suggests gene flow between them. Males from OP lineages thus likely transmit their op allele to new genomic backgrounds. This “contagious asexuality”, combined with environment-induced selection (each reproductive mode being favored under different climates) probably contributes to the long-term persistence of the *cp* and *op* alleles.

**Significance:** Asexual taxa occur in all major clades of Eukaryotes and derive from related sexual species. Yet, the genetic basis for these transitions is poorly known because crosses cannot generally be performed to genetically map the ability to propagate asexually. As a result, only one gene responsible for sex loss has been identified in one animal species. Here, using pooled genome sequencing, we identified an 840kb region (carrying 32 genes) that controls the transition to permanent asexuality in the pea aphid. We also revealed that sexual and asexual alleles diverged 0.25 million years ago and that asexual lineages likely persist through contagious asexuality, providing new insights into the mechanisms of coexistence of sexual and asexual lineages.

## Introduction

The prevalence of sexual reproduction in eukaryotes (Bell 1982) has long been considered as an evolutionary paradox, because sexual organisms transmit their genetic information twice less efficiently as asexual organisms do (Maynard Smith 1971). There is now a consensus that sex is favored over asexuality in the long-term because it purges deleterious mutations that otherwise accumulate in asexual genomes, combines favorable mutations into genomes faster and generates genotypic diversity fuelling adaptation (Muller 1964; Barton and Charlesworth 1998). Indeed, only few ancient asexual lineages exist (e.g., Mark Welch and Meselson 2000; Martens, et al. 2003), indicating the inability of asexual lineages to persist over long evolutionary time due to long-term costs. However, how sex is maintained on the short term when sexual and asexual lineages coexist is still under debate (Hartfield and Keightley 2012). The loss of sexual reproduction is observed in many animal taxa such as squamates, fishes, insects, crustaceans, nematodes and molluscs (Vrijenhoek, et al. 1989; Schon, et al. 2009). These frequent transitions from sexual to asexual reproduction reflect well the theoretical demographic advantage of asexual lineages over their sexual counterparts, which may allow them to persist over ecological times.

Sex may be lost by different ways (including interspecific hybridization, microorganism infection, spontaneous mutation or spread of contagious asexuality elements) and at various frequency, affecting the genetic features of the derived asexual lineages (Simon, et al. 2003; van der Kooi and Schwander 2014). However, little is known about the genes underlying the shifts to asexuality. Indeed, one cannot use standard crossing techniques to genetically map the ability to propagate asexually (Neiman, et al. 2014). Remarkably, certain species present lineages that only partially lost sexual reproduction, allowing the identification of the genetic bases of sex loss using recombination-based approaches. Such crosses have revealed that the genetic mechanism responsible for the transitions from cyclical to obligate parthenogenesis in aphids (Dedryver, et al. 2013; Jaquiéry, et al. 2014), rotifers (Stelzer, et al. 2010) and cladocerans (Lynch, et al. 2008; Tucker, et al. 2013; Xu, et al. 2015), and from arrhenotoky to thelytoky in hymenopterans (Lattorff, et al. 2005; 2007; Sandrock and Vorburger 2011; Aumer, et al. 2017; 2019) involves one or few loci only. However, in most cases, the precise location as well as the nature and function of the genetic determinants of these shifts to obligate asexuality remain unknown.

In animals, the alleles responsible for the loss of sex have been identified only in the Cape honeybee (Aumer, et al. 2019). Queens (and workers under particular conditions) produce haploid males by arrhenotokous parthenogenesis. However, some workers in the Cape honeybee are able to produce diploid eggs by thelytokous parthenogenesis. A single SNP in the gene *th-LOC409096* is associated with thelytokous reproduction in workers (Aumer, et al. 2019). The *th* gene encodes a receptor protein (with a transmembrane helix and a signal peptide) and locates within a non-recombining region of 60 Mb containing another gene, a hormone receptor that regulates ecdysis and juvenile hormone synthesis. The thelytoky allele (th_t_h) is dominant, contradicting previous works that suggested it was recessive (Lattorff, et al. 2005; 2007; Aumer, et al. 2017) or that thelytoky was polygenic (Chapman, et al. 2015). Hence all thelytokous individuals are heterozygous th_th_/th_ar_ and all arrhenotokous ones have the th_ar_/th_ar_genotype (Aumer, et al. 2019).

Another well studied system is *Daphnia pulex,* a crustacean reproducing by cyclical parthenogenesis, an alternation of many parthenogenetic generations and one sexual generation producing diapausing eggs, referred to as CP. In this species, sex-limited meiosis-suppressing genetic factors enable some lineages to produce diapausing eggs by parthenogenesis. These obligatory parthenogenetic lineages are called OP lineages. Genome sequencing of OP and CP lineages revealed that all *D. pulex* OP lineages share the same haplotypes in at least four genomic regions including almost two entire chromosomes and parts of two others (Lynch, et al. 2008; Tucker, et al. 2013; Xu, et al. 2015), which have been acquired by hybridization with the close species *D. pullicaria.*

The identification of candidate loci for sex loss can also shed light on the origins and evolutionary dynamics of asexual lineages and/or asexual alleles. In the Cape honeybee, the arrhenotoky haplotype (th_ar_) shows strong signs of positive selection, and seems to rescue the th_t_h allele, which is presumably associated with substantial female fitness disadvantages when homozygous, to produce thelytokous workers (Aumer, et al. 2019). This co-evolution between the two alleles likely explains why the th_th_ allele (which still provides a net fitness advantage overall) did not spread to other *Apis melifera* subspecies. In *D. pulex,* the large size of genomic regions associated with meiosis suppression in females complicates the identification of candidate genes. However, the female meiosis-suppressing factors can be transmitted by males from *D. pulex* OP lineages when they mate with females from a CP lineage, creating new OP lines by so-called “contagious asexuality”. Analyses of rates of SNP conversion between OP and CP haplotypes within lineages revealed that all OP lineages were extremely young (22 years on average, Tucker, et al. 2013). Contrastingly, the age of the radiation of the OP was much older. Based on the synonymous divergence between the different OP haplotypes, the radiation was estimated to have occurred between 1250 and 187,000 years ago, corresponding to the divergence of the OP haplotypes clade from the homologous sequences in the exclusively sexual species *D. pulicaria* (Tucker, et al. 2013). These results illustrate that, under contagious asexuality, the asexuality-conferring allele can be markedly older than OP lineages themselves. Even though each OP lineage might be doomed to extinction, the ancient asexual allele can persist by spreading in “purged” genomic backgrounds through males.

Aphids are another appropriate model for studying the genetic basis of the loss of sex. The ancestral mode of reproduction in this group is cyclical parthenogenesis, but nearly 45% of the 5,000 aphid species show partial or complete loss of sexual reproduction (Moran 1992). Typically, CP lineages undergo several successive generations of parthenogenesis (by viviparous parthenogenetic females) in spring and summer. In autumn, photoperiod shortening triggers the production oviparous sexual females and males (Le Trionnaire, et al. 2008). The winter-diapausing eggs resulting from sexual reproduction are the only frost-resistant stage of the aphid developmental cycle (Simon, et al. 2002). They give birth to viviparous parthenogenetic females in the next spring, which start a new cycle.

Interestingly, some lineages have lost the ability to produce sexual females in response to the photoperiodic cues, and thus reproduce yearlong by viviparous parthenogenesis (Simon, et al. 2002; Frantz, et al. 2006; Simon, et al. 2010). These OP lineages are demographically advantaged over CP lineages in mild winter regions, mainly because they do not undergo lengthy egg diapause. However, they cannot survive in regions with harsh winters because they are unable to produce cold-resistant eggs (Moran 1992). Thus, selection by climate results in a geographical distribution of reproductive phenotypes where OP lineages occupy regions with mild winters and CP lineages those with cold winters, both co-occurring in areas with intermediate or fluctuating climates (Rispe and Pierre 1998; Simon, et al. 2002; 2010). Interestingly, many OP lineages have retained the capacity to produce males in autumn, so that gene flow between OP and CP lineages may occur in the wild (Halkett, et al. 2008; Dedryver, et al. 2013; Jaquiéry, et al. 2014). In addition, since OP-produced males are usually fertile (Dedryver, et al. 2019), they can be crossed with CP females to identify the genetic basis of reproductive mode variation.

In the pea aphid *Acyrthosiphon pisum,* such crosses have revealed that the OP phenotype was recessive (Jaquiéry, et al. 2014). The combination of two complementary approaches – QTL mapping and low-resolution genome scan using microsatellite markers on populations submitted to divergent selection for reproductive mode – pinpointed a 10-cM genomic region located on the X chromosome controlling this trait (Jaquiéry, et al. 2014). However none of the ~24,000 scaffolds constituting the ~540-Mb pea aphid genome sequence was anchored to any of the four chromosomes (IAGC 2010) and most of the scaffolds longer than 150 kb contained assembly errors associating unlinked chromosomal regions (Jaquiery, et al. 2018). As a result, the genomic context of microsatellite markers that are linked to a focus trait could not be established. The recent release of an improved assembly of the pea aphid genome (Li, et al. 2019), in which the four largest scaffolds correspond to the four chromosomes, provides an excellent opportunity to resolve this issue.

This study aims at finely characterizing the genomic region(s) responsible of the variation of reproductive mode in the pea aphid and gaining functional and evolutionary insights into the genetic determinants of the loss of sex. To this end, we performed a high-resolution genome scan based on a pooled sequencing of 103 individuals from OP and CP populations. The improved genome assembly combined to the millions of SNP markers scattered through the genome led to the identification of an 840-kb genomic region showing strong genetic differentiation between OP and CP populations, and locating nearby the locus previously identified by Jaquiéry et al. (2014). A thorough analysis of the variants present in this region was performed to identify candidate genes underlying the variation of reproductive mode in the pea aphid and to infer the divergence time between the *op* and *cp* alleles.

## Results

### A single genomic region controls reproductive mode variation

In order to identify candidate regions involved in reproductive mode variation, we measured F_ST_ values over the whole genome (4.6 million SNPs) from pooled DNA sequencing of OP and CP populations. The average genetic differentiation between populations of different reproductive modes (OP versus CP populations) measured on the four chromosomes is low (F_ST_ = 0.024). A visual inspection of sliding windows of F_ST_ along chromosomes revealed two genomic regions of high F_ST_: a very short one (30-kb) on chromosome 1 and a longer one on the X chromosome (Figure 1A and Supplementary File 1). We found out that the short region with high Fsτ on chromosome 1 was misplaced in the v3.0 reference genome (Li, et al. 2019) and that it actually located on the X chromosome at 2 Mb from the region of highest F_ST_ (Supplementary Files 2 and 3). This short region with high F_ST_ did not meet the requirements to be considered as candidate for the control of reproductive mode variation (average F_ST_ ≥ 0.4 and more than 50 SNPs). Genomic windows that satisfied these requirements all located in an 840-kb region from position 62,895,000 to 63,735,000 of the X chromosome (Figure 1A). Remarkably, this region locates at only 750 kb from the microsatellite markers having the strongest association with reproductive mode variation (Jaquiéry, et al. 2014) (Figure 1A). This 840-kb region is highly divergent between CP and OP populations (F_ST_ = 0.349) and shows elevated differentiation in every pair of populations differing in their reproductive mode, whereas no such pattern appears for any pair of populations with the same reproductive mode (Supplementary File 4).

**Figure 1.**
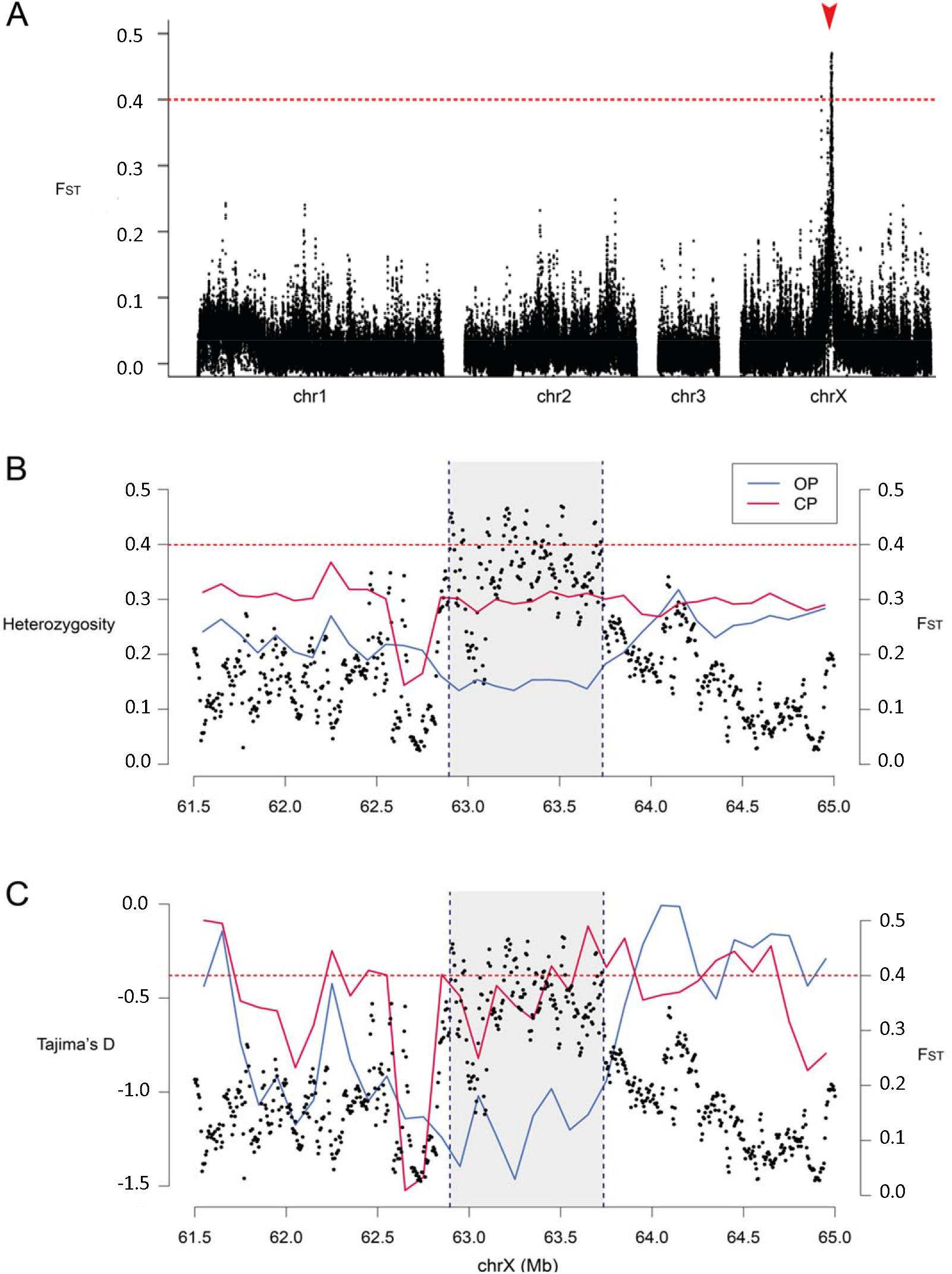
Population genetics indices computed along the A. *pisum* chromosomes. A) Genetic differentiation (F_ST_) between obligate parthenogenetic (OP) and cyclical parthenogenetic (CP) populations (20-kb windows sliding by 5-kb steps). The horizontal red dotted line represents the F_ST_ threshold at 0.4. The red arrow corresponds to the position of the outlier markers identified in Jaquiéry *et al.* (2014). Average heterozygosity (panel B) and Tajima’s D (panel C) computed per 100-kb window along the candidate region located on the X chromosome, with pink plain lines for CP populations, and blue plain lines for OP populations. Black dots represent F_ST_ values. The horizontal red dotted line represents the Fsτ threshold at 0.4 and the grey area represents the region identified as candidate for the control of reproductive mode variation.

The 840-kb candidate region contains 1,843 SNPs and 240 indels with F_ST_ > 0.5 between OP and CP populations, a value that denotes very different allele frequencies between these two population types. Heterozygosity in that region (Figure 1B) is significantly lower in OP populations (median: 0.124) than in CP populations (median: 0.302) (W = 0, p = 0.0078, two-sided Wilcoxon test using 100-kb windows as statistical units). Genome-wide heterozygosity is more similar between OP and CP populations (median values of 0.289 and 0.284 respectively) while still significantly different (W = 6507940, p < 10^-15^, two-sided Wilcoxon test using 100-kb windows as statistical units). Heterozygosity in the candidate region is lower than genome-wide heterozygosity in OP populations (U = 15.5, p = 1.01x10^-06^, two-sided Mann-Whitney test), but not significantly so in CP populations (U = 24441, p = 0.123). Median Tajima’s D values in the candidate region are also lower in OP populations than in CP populations (−1.16 and −0.48 respectively, W = 0, p = 0.0078, two-sided Wilcoxon test using 100-kb windows as statistical units), indicating the presence of a selective sweep in OP populations (Figure 1C).

We then investigated the structure of the 840-kb candidate region in the OP and CP genomes that we assembled from long-read sequences obtained from two clones (the OP X6-2 and the CP LSR1 lineage, see Supplementary File 2 for assembly quality metrics). The candidate region was located on a single scaffold on both assemblies (Figure 2A, Supplementary File 3) and did not show any large structural rearrangement between these two individual genomes. Accordingly, the sequencing depth ratio OP/(OP+CP) computed over 2-kb windows from Pool-seq data (Figure 2B) failed to reveal any large deletion in OP populations.

**Figure 2.**
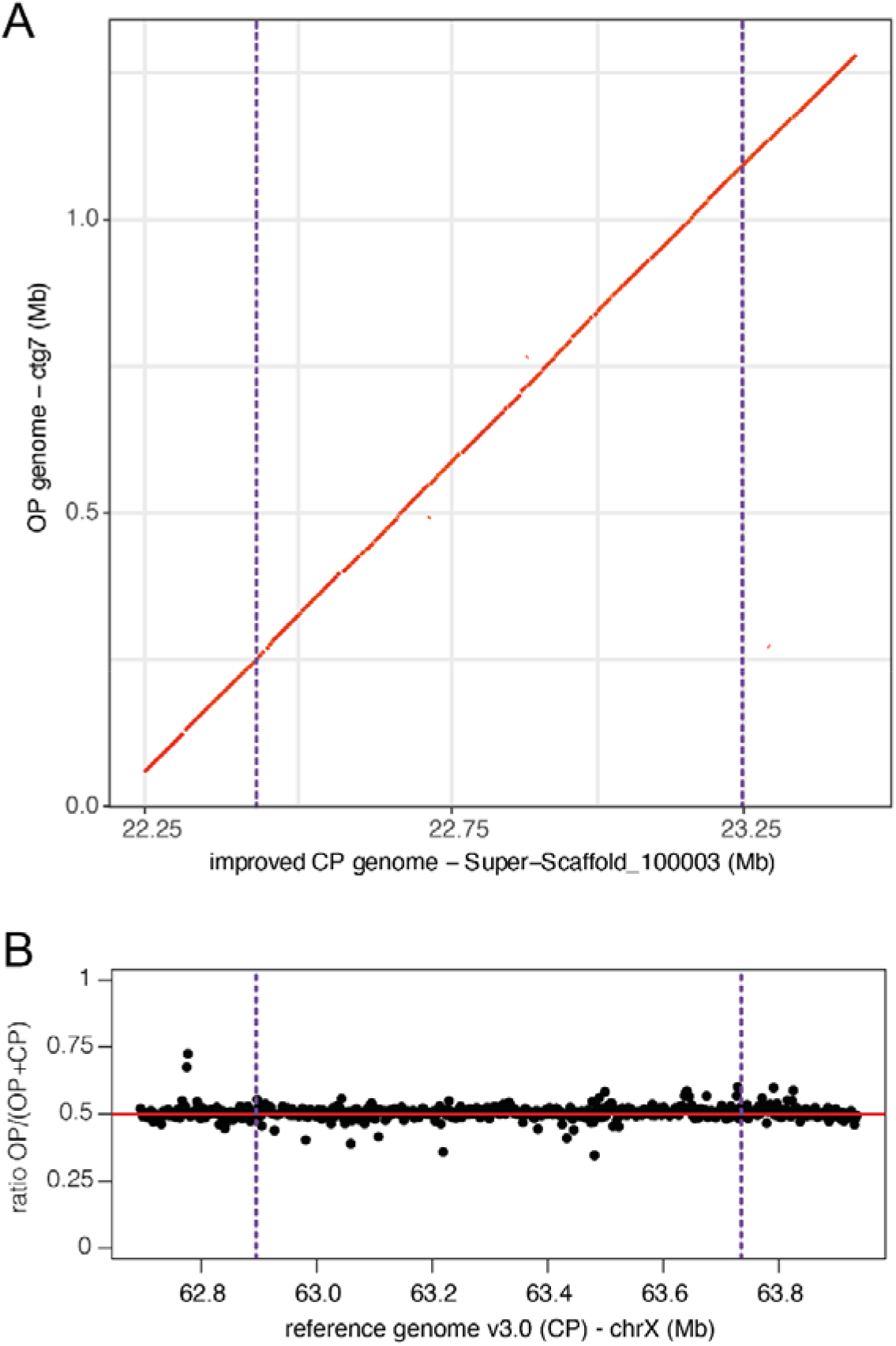
Structure of the 840-kb candidate region in obligate parthenogenetic (OP) and cyclical parthenogenetic (CP) genome assemblies. A) MUMmer alignment plot comparing parts of the two scaffolds (one per genome) containing the candidate region. B) Normalized sequencing depth ratio OP/ (OP+CP) calculated over 2-kb windows along the X chromosome. In both panels, the purple vertical dashed lines delimit the 840-kb candidate region.

### Gene content of the candidate region

The 840-kb candidate region for the control of reproductive mode contains 32 predicted genes (Table 1). Ten of these showed no homology with *Drosophila* proteins, among which nine were annotated as uncharacterized protein on NCBI and one (LOC100159148) had homologies with a nuclear pore complex protein from *Salmo trutta* (Table 1 and Supplementary File 5). The remaining 22 genes have *Drosophila* homologues, including seven that encode proteins of unknown function and 15 that are homologous to *Drosophila* genes with functional annotations and phenotypic characterizations. Interestingly, the amino acid sequences of these 15 genes all share the typical conserved protein domains identified in *Drosophila,* thus giving strong confidence in their annotation (Supplementary File 5). More precisely, four are annotated as Transcription factors, three of them sharing typical features of zinc-finger proteins (LOC1000159233, LOC100161275, LOC107882169). Seven genes are homologous to genes coding for enzymes known to be involved in general metabolism in *Drosophila:* a trimethylguanosine synthase (LOC100570687), a sphingomyelin phosphodiesterase (LOC100169137), a N-acetylglucosaminyltransferase (LOC100569179), a protein kinase (LOC100161186), a fatty acyl-coA reductase (LOC100169017), a Rho GTPase activating protein (LOC100163133) and a cysteine-type peptidase (LOC100163837). Finally, the four remaining genes are homologous to *Drosophila* genes for which phenotypic analyses of mutants revealed their involvement in key biological processes associated with germline and embryo development, including miRNA processing and RNA interference for *Cpb20* (LOC100570523) and *pasha* (LOC100168027), cell cycle control for *APC10* (LOC100165999) and dopamine signaling for *punch* (LOC100164133).

**Table 1.**
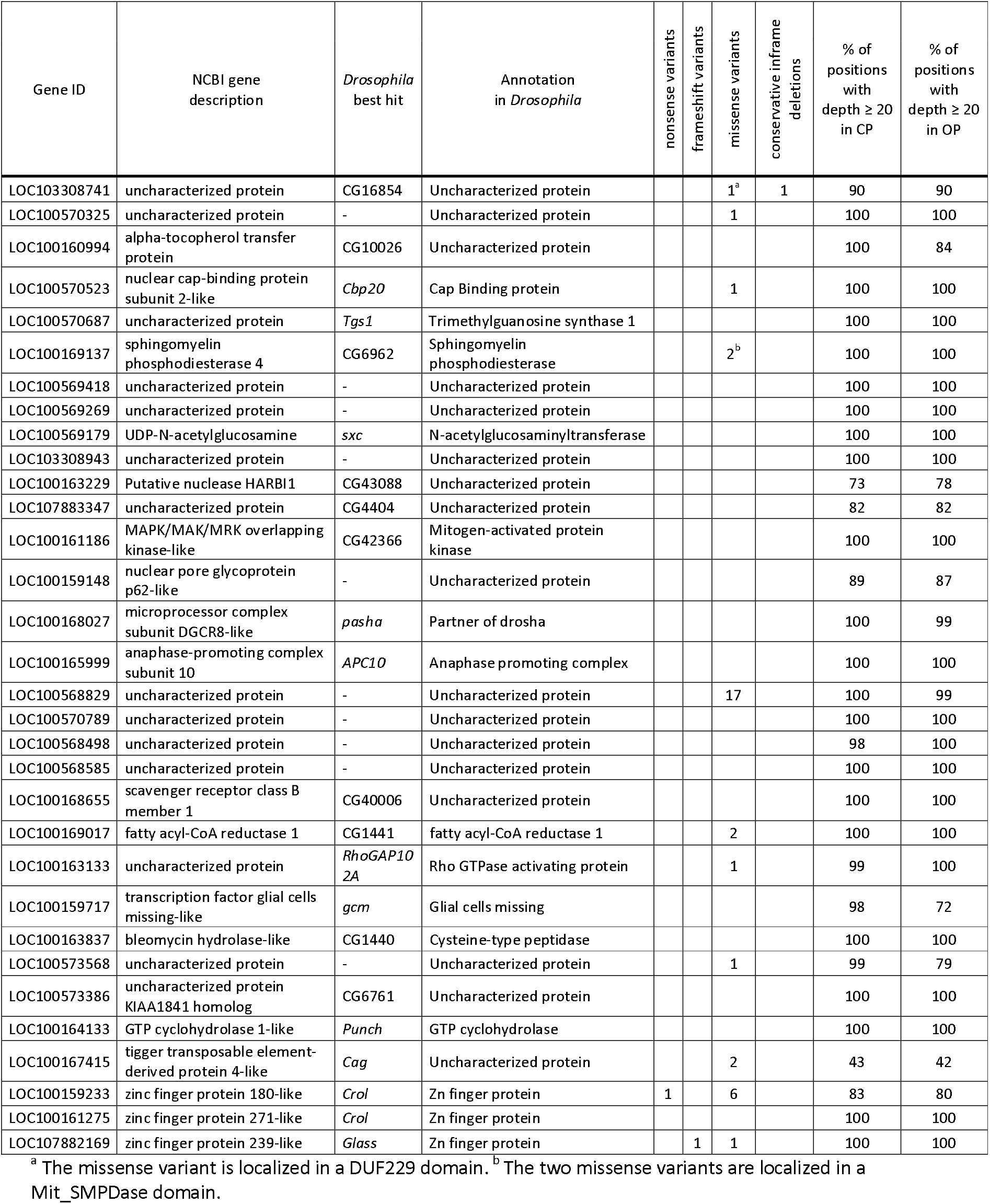
Annotation of the 32 genes predicted in the 840-kb candidate region controlling reproductive mode in pea aphids, and numbers of non-synonymous variants of different types among cyclical parthenogenetic (CP) and obligate parthenogenetic (OP) populations.

Among variants of the candidate region annotated using SnpEff, 38 impact protein sequences and show large differences in allele frequencies between OP and CP populations (F_ST_ > 0.5) (Table 1 and Supplementary File 5). These encompass 35 missense variants, one frameshift variant, one conservative in-frame indel and one nonsense variant (Table 1 and Supplementary File 5), together affecting 11 genes. Five of these are homologous to genes encoding uncharacterized proteins. Three genes with homologues in *Drosophila – Cbp20* (LOC100570523), *Fatty-acyl-CoA reductase* (LOC100169017) and *RhoGAP102A* (LOC100163133) – display one or two non-synonymous SNPs outside the typical functional domains of these proteins. Interestingly, LOC100169137 – homologous to a sphingomyelin phosphodiesterase – displays two SNPs in its mit_SMPDase domain, both changing the property of the corresponding amino acid. Finally, two genes sharing features of zinc-finger transcription factors (LOC100159233 and LOC107882169) show polymorphism possibly resulting in truncated proteins in OP lineages. The remaining 21 genes of the region do not display any polymorphism changing the protein sequence between OP and CP lineages.

There is no clear evidence for large indels associated with reproductive mode variation within the 32 genes of the candidate region, as most (29) show similar sequencing depth in OP and CP populations (Table 1 and Supplementary File 6). For each of the five genes that had less than 90% of their length sufficiently sequenced in OP and CP populations (LOC100163229, LOC107883347, LOC100159148, LOC100167415 and LOC100159233, Table 1), the same gene segment show reduced sequencing depth in both types of populations (Supplementary File 6). For three genes (LOC100160994, LOC100159717 and LOC100573568), the percentage of gene length with sufficient sequencing depth is lower in OP than in CP populations. However, this difference is supported by only one population (out of 6) in which the sequencing depth did not meet our criteria, hence failing to indicate consistent lack of coverage in all OP populations.

### Age of divergence of the op and cp alleles

To estimate the age of the divergence of the *op* and *cp* alleles, the coding sequences of the 32 genes located in the candidate region were concatenated, resulting in a 41,886-bp sequence that carries 66 variants with F_ST_ > 0.5 between OP and CP populations (Supplementary Files 7 and 8). The dS value calculated between the *op* and *cp* alleles is 0.0025, and corresponds to a divergence time of 242,504 years based on the calibrated dS between *A. pisum* and *M. persicαe.* The use of the experimentally estimated mutation rate in *A. pisum* (Fazalova and Nevado 2020) combined to the 1,843 SNPs with F_ST_ > 0.5 over the whole 840-kb region resulted in an estimated divergence time between *op* and *cp* alleles of 230,533 years (95%CI: 164,297 – 353,834).

## Discussion

In this study, we took advantage of a newly available genome assembled at the chromosomal level (Li, et al. 2019) to precisely analyse the genomic differentiation between cyclical and obligate parthenogenetic populations of the pea aphid, enabling us to pinpoint a single genomic region associated with reproductive mode variation. This 840-kb candidate region carries 32 predicted genes and harbors > 2000 SNPs and short indels that show strong differences in allelic frequencies between OP and CP populations. This reproductive polymorphism appears relatively ancient, as the *op* and *cp* alleles diverged about 0.25 million years ago. The lower heterozygosity and negative Tajima’s D values in OP populations are indicative of a selective sweep on the *op* allele, which must be favored in regions whose mild winters allow parthenogenesis all year long.

As our analysis did not reveal large structural variation between *cp* and *op* alleles, differences controlling reproductive mode probably involve only small-sized polymorphisms. SNPs and small indels are frequent along the candidate region and may affect reproductive mode by altering the function of genes controlling the switch to the sexual phase. However, none of the 32 genes located within the candidate region corresponds to those identified as differentially expressed in transcriptomic studies that investigated the photoperiod signal transduction step and the switch from sexual to asexual embryogenesis in a CP lineage (Le Trionnaire, et al. 2007; 2008; 2012; Gallot et al. 2012). These studies may have identified genes mainly acting downstream those present in the candidate region. Nonetheless, four genes of the candidate region *(cpb20, Pasha, APC10* and *punch)* share similarities with *Drosophila* genes whose functions could play a role in reproductive mode switch in aphids. Interestingly, *cpb20* is an mRNA cap binding protein involved in miRNA processing and gene silencing by RNAi: germline *Drosophila* mutants produce no eggs (Sabin, et al. 2009). *Pasha* is a double-stranded RNA-binding protein also involved in miRNA biogenesis: germline mutants do not form cysts from the germarium and fail in oocyte fate determination (Azzam, et al. 2012). *APC10* is an E3 ubiquitin-ligase that promotes metaphase to anaphase transition during the cell cycle, and germline mutants show defects in stem cells production (Liu, et al. 2016). Finally, *punch* is a GTP cyclohydroxylase involved in eye pigmentation and cell cycle control. Some mutants show defaults in dopamine synthesis and embryo development (Hsouna, et al. 2007).

Among these four genes, only *cpb20* presented a non-synonymous polymorphism with high F_ST_ between OP and CP populations. This variation lies outside the typical RNA Recognition Motif domain of the protein. Coding variation at these genes therefore appears unlikely to control the trait under scrutiny. However, five other genes of unknown function showed non-synonymous polymorphisms between OP and CP populations. A SNP and an indel in two genes containing Zinc Finger Domains are predicted to result in truncated and frameshifted products, respectively, probably leading to non-functional proteins. However, these two proteins do not share strong similarities with well-characterized *Drosophila* transcription factors; it is thus difficult to predict any phenotypic consequence of their disruption. The remaining non-synonymous polymorphisms were essentially observed in genes of unknown functions and principally located outside predicted protein functional domains.

Polymorphisms outside protein coding sequences could also control reproductive mode variation in pea aphids, as intergenic and intronic regions contain DNA motifs to which regulatory factors may bind. Transcriptomic analyses of OP and CP lineages submitted to long and short photoperiod regimes would allow testing whether some of the 32 genes are differentially expressed, thus whether they could control the reproductive polymorphism through differences in protein levels. A parallel can be drawn with *Daphnia pulex,* where male production is genetically controlled (Innes and Dunbrack 1993; Innes 1997). A recent genome scan analysis pinpointed a single gene whose male-producing and non-male producing alleles differ by seven non-synonymous substitutions (Ye, et al. 2019). These alleles are also expressed at different levels in response to the environmental cue normally inducing the production of males (Ye, et al. 2019). Whether pea aphid reproductive polymorphism is determined by expression levels, by protein variants, or by a combination of both remains an open question.

The estimated time of divergence between the *op* and *cp* alleles, at about 230,000 to 242,000 years depending on the approach used, is framed by different estimates for the age of the radiation of the pea aphid complex. These estimates vary considerably, from 18,000–47,000 years when using the divergence of the maternally inherited obligatory endosymbiont *Buchnera aphidicola* (Peccoud, et al. 2009) to 419,000–772,000 years when using nuclear divergence (Fazalova and Nevado 2020). Such wide-range variation does not permit us to determine whether the *op* allele appeared before or after the pea aphid radiation. This uncertainty could be clarified by testing whether reproductive mode in other pea aphid host races (which also present OP and CP lineages, Frantz, et al. 2006) is controlled by homologous *op* and *cp* alleles or by an independent genetic variation. However, the fact that most pea aphid host races still hybridize (Peccoud, et al. 2009; Peccoud, et al. 2014; Peccoud, et al. 2015) might make it difficult to determine whether any shared polymorphism actually emerged before the start of their divergence.

With these considerations in mind, the age of the *op* allele is likely to be much older than that of most OP lineages carrying it. As proposed by the contagious asexuality hypothesis, new OP lineages may frequently be produced by crosses between males and females carrying the *op* allele, which is recessive in the pea aphid (Jaquiéry, et al. 2014). The probability of this scenario depends on the unknown frequency of the *op* allele among CP lineages. In *Daphnia pulex,* asexual lineages are estimated to be 22 years old on average, while the asexual allele is at least 1,250 to 187,000 years old (Tucker, et al. 2013). In the pea aphid, a combination of factors probably allowed the long-term coexistence of the cp and *op* alleles. First, strong contrast in winter temperatures among western European regions allows both types of populations to stably persist in the areas where they are each adapted. Second, gene flow between OP and CP lineages may generate new OP lineages, under the scenario described above. Such gene flow is strongly supported by low genetic differentiation between OP and CP populations (genome-wide average F_ST_ of 2.4%). Mating may occur between sexual females from CP lineages and males from OP lineages, which likely coexist in regions whose lowest temperatures fluctuate around the limit tolerated by aphids. These crosses would allow the *op* allele to escape from linked deleterious mutations that may accumulate in OP lineages. They can also generate new OP lineages, ensuring the long-term persistence of OP populations (and *op* allele) through contagious asexuality. Although this scenario needs to be tested in nature for the pea aphid, previous works already showed its validity in natural populations of other aphid species (Halkett, et al. 2008).

Interestingly, two loci of considerable ecological relevance have been found on the X chromosome: the one studied here and *aphicarus,* a locus controlling the presence of wings in males (Braendle, et al. 2005; Li, et al. 2020). Remarkably, the two loci are physically close, at positions 62,56-62,75 Mb for *aphicarus* and 62,89-63,73 Mb for the one controlling reproductive mode variation. We observed a specific genetic signature at the location of *aphicarus* (Figure 1), which includes a drop of F_ST_ between OP and CP populations, and a relative drop of both heterozygosity and Tajima’s D in CP populations. The frequency of the wing-inducing allele is low in the alfalfa-adapted host race in Europe (around 5-10%, Frantz, et al. 2009; Li, et al. 2020), which could explain low differentiation between OP and CP populations and the low Tajima’s D. Indeed, the presence of only 5-10% of the highly divergent winged allele could inflate the number of low-frequency polymorphisms, hence decrease Tajima’s D.

To conclude, this work refines the size, location and gene content of the locus controlling reproductive mode in the pea aphid. Further functional studies are needed to identify the gene(s) driving reproductive mode variation, and to determine whether variation at this trait depends on variation in protein sequence and/or protein levels. Transcriptomic analyses of OP and CP lineages submitted to long and short photoperiod regimes should help to identify the causal gene(s) and underlying mechanisms. CRISPR/Cas9 targeted mutagenesis, which has been successfully developed in the pea aphid (Le Trionnaire, et al. 2019), would then allow a functional validation of the role of candidate genes. Furthermore, exploring the genetic basis of sex loss in other host races and species should clarify whether reproductive mode variation, which is widespread in aphids (Moran 1992; Simon, et al. 2002), relies on common or independent mechanisms. Finally, sequencing individual OP lineages in various populations would allow assessing the accumulation of deleterious mutations in these clones and whether climatic or other environmental factors primarily dictate their fate.

## Materials and Methods

### Aphid sampling

This study is based on the A. *pisum* samples previously used to conduct a low-density microsatellite-based genome scan (Jaquiéry, et al. 2014). Briefly, parthenogenetic females were collected on *Medicαgo sαtivα* in alfalfa cultivated fields from six sampling sites (Supplementary File 9). Three sites locate in north-east France and Switzerland, where only CP lineages can survive cold winters. The three other sites locate in south-west France where winters are generally mild and therefore favor obligate parthenogenesis (OP). For each of the six geographical populations, we succeeded to keep alive 14 to 21 genetically distinct clonal lineages, each initiated by a sampled female (Supplementary File 9).

### Pool sequencing

DNA was extracted from four fourth instar larvae per clonal lineage using the Qiagen DNeasy Blood and Tissue Kit (Qiagen, Hilden, Germany) following the manufacturer instructions. After RNAse treatment, DNA solutions were pooled in equimolar proportions for each population ensuring that clonal lineages contributed equivalent amounts of DNA to the pool. Two independent paired-end libraries were constructed per population from these DNA pools using the Genomic DNA Sample Preparation Kit (Illumina, San Diego, CA) (technical replicates). The resulting 12 libraries were sequenced on four lanes of the Illumina HiSeq 2000 platform in a single 2×100-cycle run using Illumina Sequencing Kit v3. The raw data are publicly available at the Sequence Read Archive of the NCBI database, under the BioProject ID PRJNA454786.

### Mapping

For read mapping enabling variant calling, we used the v3.0 reference genome of the pea aphid (NCBI: pea_aphid_22Mar2018_4r6ur, Li, et al. 2019). This assembly is 541 Mb in size and consists of four main scaffolds corresponding to the three autosomes and the X chromosome, and 21,915 additional short scaffolds not positioned on chromosomes (which account for 14% of the bases, Li, et al. 2019). Paired-end reads were mapped to a fasta file containing the *A. pisum* reference genome v3.0 and the sequences of its known endosymbionts (Guyomar, et al. 2018) with bwa-mem v0.7.10 (Li and Durbin 2009), using defaults parameters. Low quality (Q<20) and improperly paired alignments were removed with SAMtools v1.2 (Li, et al. 2009). Read pairs corresponding to duplicates were then identified with Picard Markduplicates v2.18.2 (http://broadinshtute.github.io/picard/) and removed.

### Variant calling

The 12 alignment (BAM) files corresponding to the 12 DNA libraries were merged in a single mpileup file using SAMtools (Li, et al. 2009) and a sync file was created using Popoolation2 (Kofler, et al. 2011) with default parameters except for a minimum base quality set to 20. Positions corresponding to the aphid symbionts and mitochondria were removed from the sync file, to analyse the pea aphid nuclear genome only. The sequencing depth per library ranged from 15.1 to 20.4x (Supplementary File 9).

Stringent filters were applied to select reliable and informative SNPs for the genome scan. Firstly, only biallelic SNPs for which each allele was carried by at least four reads, considering all libraries, were kept. If three alleles were scored at a position, including one representing only one read or a deletion, this allele was ignored and the SNP was considered biallelic. Secondly, only SNP positions with a sequencing depth higher than 20 and lower than 60 per population were considered, the mean depth ranging from 31 to 37 depending on the population. The upper limit of 60 was chosen to avoid duplicated genomic regions not resolved in the reference genome, and the lower limit of 20 to discard SNPs whose sampling was too low for reliable allele frequency estimates. Thirdly, a minor allele frequency threshold of 5% was applied to eliminate SNPs harboring rare alleles and which are not informative for a genome scan. After applying these selection criteria, we obtained a dataset of 4,633,747 SNPs, 94.7% of these SNPs locating on the four largest scaffolds corresponding to the four chromosomes.

### Detection of genomic regions associated with reproductive mode variation

To visualize the structure of the dataset, a Principal Component Analysis (PCA) was carried out on the 12 libraries with prcomp, implemented in R version 3.6.1 (R Core Team 2019), using their allele frequencies at 50,000 randomly sampled SNPs. Since the two libraries from each population grouped together (Supplementary File 10), we summed their allele counts as if they constituted only one library. To estimate differentiation between types of populations with different reproductive modes, we also summed allele counts for the three CP populations on one hand and for the three OP populations on the other hand. The summed allele counts were used to calculate F_ST_ at each SNP between reproductive modes with the R package poolfstat, which implements F_ST_ estimates for Pool-seq data (Hivert, et al. 2018). We then calculated the average F_ST_ within 20-kb windows sliding by 5-kb steps to smooth its variation along the genome. To select regions of elevated genetic differentiation between population types, we considered windows characterized by average F_ST_ values ≥ 0.4 and with more than 50 SNPs, the average number of SNPs per window being close to 120. F_ST_ were computed similarly for each pair of populations.

Heterozygosity (H_e_, following Nei 1973) was calculated per type of populations (OP or CP) at each SNP using allele frequencies derived from allele counts. The mean H_e_ per population type was then computed in 100-kb contiguous windows. Finally, to detect selective sweeps resulting from selection on reproductive mode variation, Tajima’s D was calculated for each population type. For this, we used the pileup-formatted SNP files for the pool samples of each reproductive mode (*i.e*., one OP and one CP population) generated previously. We randomly subsampled the datasets as recommended to achieve a uniform depth using PoPoolation 1.2.2 (Kofler, et al. 2011), using the following parameters: --target-coverage 72 -max-coverage 360 -min-qual 20. Tajima’s D were then calculated using PoPoolation 1.2.2 (Kofler, et al. 2011) over 100-kb non-overlapping windows with the following parameters: -min-count 2 -min-covered-fraction 0.5.

### Gene and variant annotation

The above analyses identified a single genomic region as candidate for the control of reproductive mode variation. Amino acid sequences of the predicted genes present in this region were retrieved from the v3.0 version of the pea aphid genome assembly. Annotations for these genes were obtained from the general feature format (gff) file available on NCBI (GCF_005508785.l_pea_aphid_22Mar2018_4r6ur_genomic.gff.gz). Whenever a gene had multiple predicted transcripts, we only kept the longest transcript (Supplementary File 5). A BlastP analysis (Altschul, et al. 1990) was then performed against Flybase (http://flybase.org/) to identify the closest *Drosophila* homologue for each of these aphid genes (at p<10^-7^). Conserved protein domains were identified and annotated for each gene using the SMART web resources (http://smart.embl-heidelberg.de/; Letunic, et al. 2020) with the “normal” mode and a significance level of 10^-10^. We examined the variants (SNPs and short indels) included in the candidate region for reproductive mode variation to detect potentially causal polymorphisms. These variants were classified according to their impact on gene structure by SnpEff v4.3t (Cingolani, et al. 2012) with default parameters and using the GFF file available on NCBI. Variants with moderate to high predicted impact were retained for further analysis. This includes variants resulting in premature stop codons, frameshifts, missenses or conservative in-frame indels. When not already available from our previous calculations, F_ST_ were calculated between the OP and CP populations for each of these variable positions, to assess its correlation with reproductive mode. For this computation, we retained only positions with a sequencing depth ≥ 20 in every population.

### Comparison of the structure of the candidate region in CP and OP genomes

To compare the structure of the candidate region between CP and OP genomes, we assembled the genome of an OP lineage (clone X6-2, Jaquiéry, et al. 2014), as the A. *pisum* reference genome v3.0 (Li, et al. 2019) was assembled from a CP lineage (clone LSR1, IAGC 2010). Oxford Nanopore technology was used to obtain long-read sequences from the OP lineage and to build a *de novo* genome assembly (see Supplementary File 2 for details). We also found that the A. *pisum* reference genome v3.0 (Li, et al. 2019) contained some small assembly errors which could impact our results (see results and Supplementary Files 2 and 3). We therefore constructed a new assembly for the LSR1-CP lineage (referred to as “improved CP genome” hereafter) with ONT- and PacBio-generated long reads and optical map data (Supplementary File 2). We then compared the structure of genomes assemblies at a 1.25-Mb region containing the 840-kb candidate region using MUMmer v3.22 (Kurtz, et al. 2004). Pairwise alignments of the CP and OP genome sequences were assessed using NUCmer v3.07. Results were filtered using the delta-filter script to keep optimal correspondence with a minimum length of 1000 bp and a minimum alignment identity of 90%, and were visualized using MUMmerplot v3.5 (Kurtz, et al. 2004). Complementarily, to investigate the deletion of short genomic regions in OP populations, we plotted the sequencing depth ratio OP / (OP+CP) from the Pool-seq data. The sequencing depths of the OP and CP populations were normalized prior to ratio calculation, so that a ratio of 0.5 is expected for genome segments presenting the same copy number in the OP and CP populations. To visualize results, we computed the average of this ratio over 2-kb non-overlapping windows on the candidate region.

### Age of divergence of op and cp alleles

To estimate the divergence time between the *op* and *cp* alleles of the candidate region, *op* and *cp* consensus sequences were established from SNPs showing F_ST_ value > 0.5 between CP and OP populations (which corresponds to allele frequency differences typically greater than 0.55 for a biallelic SNP) and a sequencing depth ≥ 20 in every population (Supplementary Files 7 and 8). We analysed these sequences by two different dating approaches. The first relied on synonymous mutation rates (dS). dS was computed between concatenated *op* and *cp* coding sequences (extracted from the consensuses) with the seqinr R package (Charif and Lobry 2007). We then assumed that the synonymous mutation rate per time unit between *cp* and *op* alleles was the same as that between the pea aphid and the peach-potato aphid *Myzus persicαe,* whose divergence is estimated at some 22 million years ago and corresponds to a dS of 0.2268 (Johnson, et al. 2018; Mathers, et al. 2020).

The second approach used the full *op* and *cp* consensus sequences and the per-nucleotide mutation rate that has been estimated in the pea aphid as *μ*_parth_ = 2.7×10^-10^ (95% CI: 1.9×10^-10^ - 3.5×10^-10^) per parthenogenetic generation from a mutation accumulation experiment (Fazalova and Nevado 2020). The annual mutation rate for an OP lineage was thus estimated as *μ*_op_ = *N*_gen_ x *μ*_path_ *N_sen_* being the number of generations per year (estimated at 15). For a CP lineage, we followed Fazalova and Nevado (2020) and estimated the mutation rate as *μ*_cp_ = (*N*_gen_-l) × *μ*_parth_ + *μ*_se_, where *μ*_sex_ (2.96×10^-9^; 95% CI: 1.52×10^-9^-4.99×10^-9^) is the average mutation rate per sexual generation in insects (Keightley, et al. 2014; Keightley, et al. 2015; Yang, et al. 2015; Liu, et al. 2017; Oppold and Pfenninger 2017) as there is no such estimate for aphids. The time of divergence (*T*) between the *op* and the *cp* alleles was then estimated as *T* = *N*_mutatedsites_/(2 ×*N*_sites_×*μ*), where *μ* = (*μ*_cp_+ *μ*_cp_)/2. *N*_mutated_ is the number of SNPs with F_ST_ > 0.5 in the 840-kb candidate region (1,843), and *N*_sites_ the number of sites with sequencing depth ≥20 in every population (740,918).

## Acknowledgments

This work was supported by grants from the French Research Agency (SexAphid ANR-09-GENM-017-001 and Speciaphid ANR-11-BSV7-0005), the INRAE-SPE Department (AAP GenAsex and half a PhD grant for HD and PN), the Region Bretagne (ARED, half a PhD grant for HD and PN) and the European Union’s Horizon 2020 research and innovation programme under the Marie Skłodowska-Curie Action (grant agreement no. 764840 for the ITN IGNITE project).

## Data Availability

Raw sequence reads are deposited in on NCBI (PRJNA454786 and PRJNA745262). Genome assemblies will be available upon article acceptance at the following permanent addresses: https://bipaa.genouest.org/sp/acyrthosiphon_pisum/download/genome/LSR1_CP https://bipaa.genouest.org/sp/acyrthosiphon_pisum/download/genome/OP

