## Supplementary Files 1-10 for "A quarter-million-year-old polymorphism drives reproductive mode variation in the pea aphid"

**Supplementary materials**

**Supplementary File 1.** Mean F_ST_ values in 20-kb windows sliding along the *A. pisum* v3.0 reference genome by 5-kb steps. The red arrow indicates a short misassembled high-Fst region on chromosome 1, which actually locates on the X chromosome (see Supplementary Files 2 and 3 for details). The inset corresponds to an enlargement of this region, with each point corresponding to a 1-kb window.

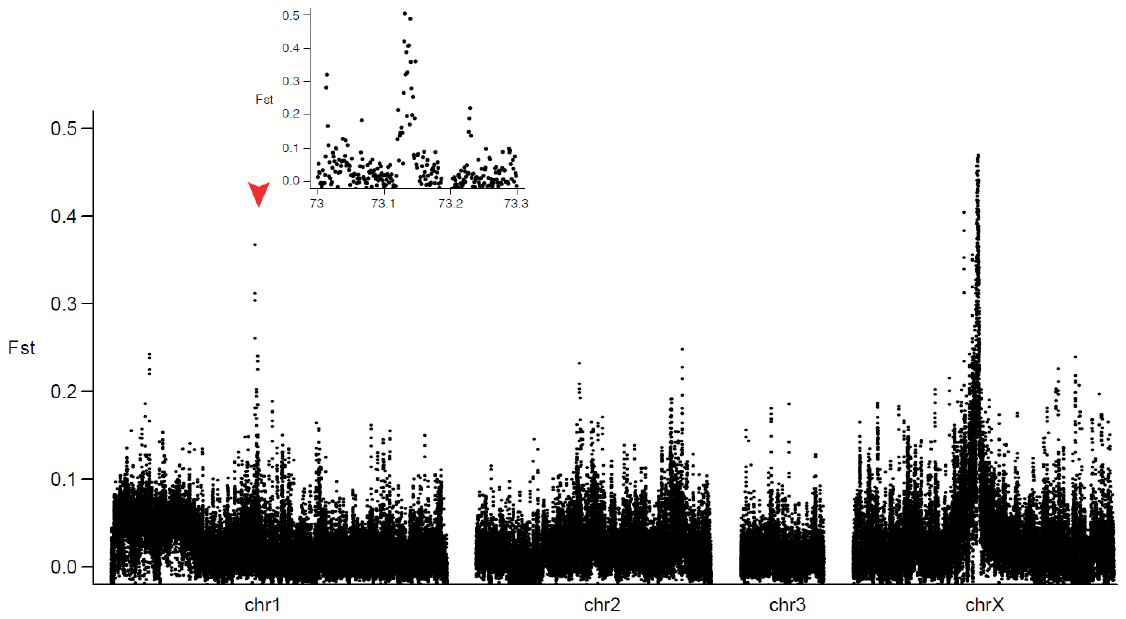

**Supplementary File 2.** This supplementary file describes the new data that were sequenced to build new genome assemblies for OP and CP lineages of the pea aphid, the construction of these assemblies and the analyses that were performed on the 840-kb candidate region with these genomes.

*OP genome sequencing and assembly*

We sequenced and assembled an OP genome with long reads (Oxford Nanopore, ONT). The lineage selected for sequencing (lineage X6-2 from cross 6 in the Supplementary Figure S1 from Jaquiéry, et al. 2014) was characterized as OP in conditions inducing sexual morph production. DNA was extracted from parthenogenetic embryos dissected from 20 adult females of this X6-2 OP lineage. Embryos were grounded in lysis buffer using a potter. The lysis solution was then used in a Phenol protocol using high vacuum grease silicone to separate aqueous and organic phases without pipetting, and low centrifuge speed to preserve DNA integrity. The DNA was directly precipitated with isopropanol and then resuspended in ultra-pure sterilized water. DNA concentration was measured with Qubit fluorometer and DNA fragment length (range 17kb-100kb) was assessed with both a Pulse Field Gel Electrophoresis and a Femto Pulse Run.

A Nanopore genomic DNA library was then prepared using the Ligation Sequencing Kit (Oxford Nanopore Technologies), following the manufacturers protocol, and sequenced on an R9.4 flow cell for 72 hours (ONT GridION technology). See Table 1 (in the present file) for a summary of the characteristics of sequence data obtained, which are publicly available on NCBI (BioProject ID PRJNA745262).

The Nanopore reads were first trimmed with PoreChop v0.2 (Wick, et al. 2017), using default parameters. Reads were then assembled with wtdbg v2.5 (Ruan and Li 2020) with the options -x ont -g 500m and polished with the same tools after an alignment step of the trimmed reads with minimap v2.14 (Li 2018). The characteristics of this OP genome assembly are summarized in Table 2 (this file). This genome assembly will be publicly available at <https://bipaa.genouest.org/sp/acyrthosiphon_pisum/download/genome/OP> after article publication.

*Improved CP genome sequencing and assembly*

As the genome available for a CP lineage of the pea aphid (reference v3.0, Li, et al. 2019) contains small assembly errors – which could have affected our conclusions if the candidate region was involved – we chose to build a new genome assembly for a CP lineage. To do so, we combined PacBio and ONT sequencing data with an optical map performed on a CP lineage of the pea aphid (clone LSR1).

For the optical map, ultra-high molecular weight (uHMW) DNA was purified from 0.2 g of frozen larvae from the LSR1 aphid lineage according to the Bionano Animal Tissue DNA Isolation Grinding Protocol (800002 - Bionano Genomics) with the following specifications and modifications. Briefly, the aphid larvae were disrupted in the homogenization buffer with a potter. Nuclei were washed and then embedded in agarose plugs. After overnight proteinase K digestion in the presence of Lysis Buffer (Bionano Genomics) and one hour treatment with RNAse A (Qiagen), plugs were washed four times in 1x Wash Buffer (Bionano Genomics) and five times in 1x TE Buffer (ThermoFisher Scientific). Then, plugs were melted two minutes at 70°C and solubilized with 2 µL of 0.5 U/µL AGARase enzyme (ThermoFisher Scientific) for 45 minutes at 43°C. A dialysis step was performed in 1x TE Buffer (ThermoFisher Scientific) for 45 minutes to purify DNA from any residues. The DNA samples were quantified using the Qubit dsDNA BR Assay (Invitrogen). Megabase-sized DNA fragments were visualized by pulsed field gel electrophoresis (PFGE).

Labeling and staining of the uHMW DNA were performed according to the Bionano Prep Direct Label and Stain (DLS) protocol (30206 - Bionano Genomics). Briefly, labeling was performed by incubating 750 ng genomic DNA with 1× DLE-1 Enzyme (Bionano Genomics) for two hours in the presence of 1× DL-Green (Bionano Genomics) and 1× DLE-1 Buffer (Bionano Genomics). Following proteinase K digestion and DL-Green cleanup, the DNA backbone was stained by mixing the labeled DNA with DNA Stain solution (Bionano Genomics) in presence of 1× Flow Buffer (Bionano Genomics) and 1× DTT (Bionano Genomics), and incubated overnight at room temperature. The DLS DNA concentration was measured with the Qubit dsDNA HS Assay (Invitrogen).

Labelled and stained DNA was loaded on the Saphyr chip. Loading of the chip and running of the Bionano Genomics Saphyr System were all performed according to the Saphyr System User Guide (30247 - Bionano Genomics). Data processing was performed using the Bionano Genomics Access software (<https://bionanogenomics.com/support-page/bionano-access-software/>). A total of 1.1 Tb data were generated. From this data, molecules with a size larger than 150 kb, the threshold for map assembly, represent 356 Gb data. These filtered data (> 150 kb), corresponding to 651x coverage of the 550 Mb estimated size of *A. pisum* genome, were compiled from 1,557,530 molecules with N50 of 223 kb and an average label density of 13.2/100kb. The filtered molecules were aligned using RefAligner with default parameters. It produced 127 genome maps with a N50 of 20 Mb for a total genome map length of 778.5 Mb. As the map size was longer than expected, due to the heterozygosity, we purged it in order to obtain only one haplotype for each optical map before the hybrid scaffolding step. For that, we first used runCharacterise from Bionano tools to align maps with each other, and created an alignment file (xmap file). From that file, we recovered supernumerary maps which align globally to other maps, with an in-house java program. This way, we purged the 778.5 Mb optical maps and obtained a genome map length of 527 Mb (consistent with the pea aphid genome size), consisting in 34 maps.

For PacBio sequencing, high molecular weight DNA was extracted from the CP lineage (LSR1) following a protocol similar to the one used for ONT sequencing of the OP clone (see above). PacBio genomic DNA libraries were prepared using the SMRTbell Template Prep Kit 1.0 (PacificBiosciences) following the manufacturer’s protocol and sequenced with six SMRTCells 1M on PacBio Sequel (PacificBiosciences) in Gentyane Platform (Clermont-Ferrand, France) and Centre for Genomic Research (University of Liverpool, UK). The raw data (see Table 1 below) are publicly available on NCBI (BioProject ID PRJNA745262). PacBio raw reads were first treated to produced CCS with the ccs program of the suite PacificBioSystems Pitchfork v3.0 (commit 96f0b06, https://github.com/PacificBiosciences/pitchfork) with the option --maxLength 20000. The resulting CCS and subreads from reads with no CCS were mixed. ONT sequencing was also performed on the LSR1 CP lineage, using the same protocol as described above for the OP lineage.

ONT reads and PacBio subreads were then aligned to the *Buchnera aphidicola* strain APS complete genome (NC_002528.1) with minimap2 v2.17 (Li 2018) with default parameters. The reads matching *B. aphidicola* genome were removed for further analyses. The final set of reads was then assembled and polished with flye v2.7.1 (Kolmogorov, et al. 2019) following the website instructions (https://github.com/fenderglass/Flye) for a mix of ONT and PacBio sequences (assembly of the 2 sets with the options -g 530m - iterations 0, then polishing with PacBio data only with the options -resume from polishing - genome-size 530m).

Then, the 34 reduced optical maps were compared individually to the genome sequences with the hybridScaffold.pl script from Solve3.6.1_11162020 (with the options -B 2 and -N 2), and finally all the sequences resulting of hybrid maps were merged into one genome sequence. Lastly this genome sequence was gapfilled with LR_Gapcloser v1.1 (Xu, et al. 2018) (with the option -s p) and the reads corrected with CANU v1.9 (Koren, et al. 2017).

We refer to this final assembly as the “improved CP assembly” (see Table 2 below for its main characteristics), which will be available at <https://bipaa.genouest.org/sp/acyrthosiphon_pisum/download/genome/LSR1_CP> after article publication.

**Table 1.** Sequencing data used for genome assemblies

|  | **PacBio data** | **ONT (Nanopore)** | |
| --- | --- | --- | --- |
| Aphid lineage | LSR1 (cyclically parthenogenetic) | LSR1 (cyclically parthenogenetic) | X6_2 (obligate parthenogenetic) |
| Number of reads | 8,007,204 | 372,636 | 1,330,350 |
| Sequenced bases | 51,786,985,914 | 4,240,655,565 | 15,864,369,067 |
| Genome coverage | 97.7X | 8X | 30X |
| Mean read length (bp) | 6,468 | 11,380 | 11,924 |
| Median read length (bp) | 6,163 | 9,010 | 10,262 |
| N50 (bp) | 8,750 | 17,775 | 16,988 |

**Table 2.** Genome assembly statistics. Busco analyses were realized with BUSCO 4.0.6, using the dataset insecta_odb10 (1367 BUSCOs, https://busco.ezlab.org/list_of_lineages.html).

|  | **Improved CP genome** | **OP genome** |
| --- | --- | --- |
| Aphid lineage | LSR1 (cyclically parthenogenetic) | X6_2 (obligate parthenogenetic) |
| Data used | PacBio, Nanopore, Optical map | Nanopore |
| Number of scaffolds | 32 | 3889 |
| Assembly size (Mbp) | 527 | 478 |
| N50 (Mbp) | 60 | 0.955 |
| L50 | 4 | 127 |
| BUSCO - single | 1220 | 902 |
| BUSCO - Duplicated | 63 | 8 |
| BUSCO - fragmented | 13 | 186 |
| BUSCO - Missing | 71 | 271 |

*Identification of the actual chromosomal localization of the 30-kb outlier region found on chromosome 1 on the v3.0 reference genome*

Our genome scan identified two main genomic regions with high F_ST_ values, the main 840-kb X-linked candidate region and a short one (30 kb) on chromosome 1 (located between 73,118,603 and 73,151,851, see Supplementary File 1). Given the abrupt changes in F_ST_ at the border of this 30-kb region, we suspected a genome assembly error. In a previous study, the entire *A. pisum* genome was assigned to the X or autosomes based on ratios of sequencing depth in males (X0) to females (XX) (Jaquiéry, et al. 2018). Using these data, we discovered that the 30-kb region was indeed misplaced on the v3.0 reference genome (Li, et al. 2019) and actually belongs to the X chromosome. We located the 30-kb chr1 region on Super-Scaffold_100003 of the improved CP genome, between positions 20,170,923 and 20,205,127 (see Supplementary File 3A). Super-Scaffold_100003 corresponds to a region on the X chromosome of the v3.0 reference genome (Li, et al. 2019) and the 30-kb region would be localized between positions 60,702,513 and 60,707,859. This places the short region of high F_ST_ at ~2 Mb of the main 840-kb candidate region (which also locates on Super-Scaffold_100003, see Supplementary File 3A). These regions may thus be under the influence of the same locus controlling reproductive mode. However, since the F_ST_ values of this 30-kb region were below the F_ST_ threshold of 0.4, we did not further retain it as a candidate region.

*Structure of the 840-kb candidate region in OP and CP genomes*

Finally, to characterize the genomic structure of the 840-kb candidate region, we located this region in the OP genome and in the improved CP genome. In the improved CP genome, we located the 840-kb candidate region on the Super-Scaffold_100003, between positions 22,415,764 and 23,247,663 (Supplementary File 3A). In the OP genome, the whole 840-kb candidate region was also found on a single scaffold (between positions 249,000 and 1,093,000 on scaffold cgt7). Pairwise alignments of the sequence corresponding to the 840-kb region (flanked by 200 kb on each side) in the 3 different genome assemblies (the v3.0 reference genome, the improved CP genome and the OP genome) were assessed using NUCmer v3.07 from package MUMmer v3.22 (Kurtz, et al. 2004). Alignments were filtered using the script delta-filter to keep optimal correspondence with a minimum length of 1000 bp and a minimum alignment identity of 90. We found that the local assembly of the region corresponding to the 840-kb region was concordant between the improved CP assembly and the v3.0 reference assembly and also with the OP genome assembly (Supplementary File 3B and 3C, see also Figure 2 in the manuscript).

**Supplementary File 3.** Panel A: location of the 30-kb chromosome 1 region and of the 840-kb candidate region on Super-Scaffold_100003 from the improved CP genome assembly. Panels B and C: MUMmer alignment plots for the nucleotide sequences from the different genome assemblies corresponding to the 840-kb candidate region (plus 200 kb on each side). The x-axis represents coordinates of the v3.0 reference genome assembly. The y-axis represents coordinates of Super-Scaffold_100003 from the improved CP genome assembly (panel B) or of scaffold ctg7 from the OP genome assembly (panel C). The purple vertical dashed lines delimit the 840-kb candidate region.

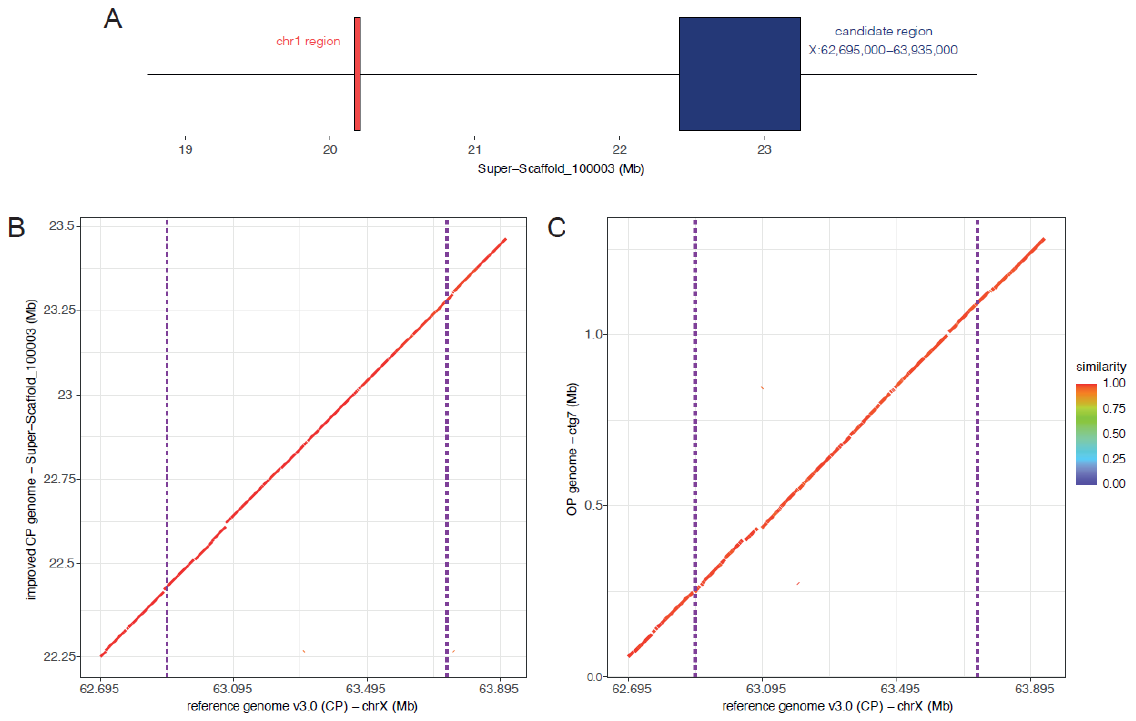

**Supplementary File 4.** Proportion of SNPs with F_ST_ values >0.5 in 20-kb windows sliding along the X chromosome by 5-kb steps, for each population pair. Panel A: populations presenting the same reproductive mode; panel B: populations presenting different reproductive modes. OP population names are written in blue and CP population names in red.

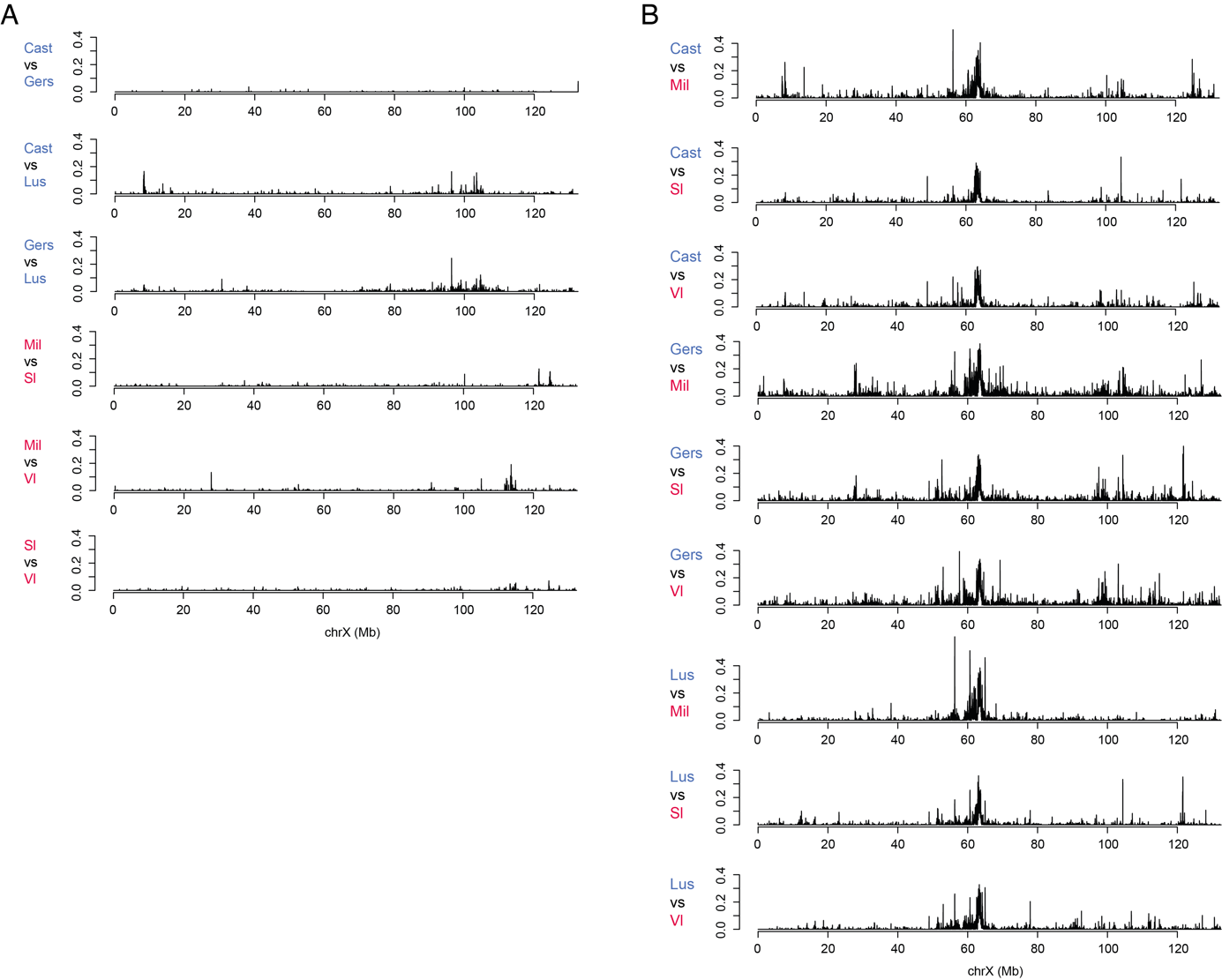

**Supplementary File 5.** Description and annotation of the 32 genes and of the variants with moderate to high impact in the 840-kb candidate region.

*See attached Excel file SupplementaryFile5.xlsx.*

**Supplementary File 6.** Summary of the coverage of the 32 genes located in the candidate region in OP and CP populations

*See attached Excel file SupplementaryFile6.xlsx.*

**Supplementary File 7.** List of the SNPs from the 840-kb candidate region with F_ST_ > 0.5 between CP and OP populations.

*See the attached Excel file SupplementaryFile7.xlsx.*

**Supplementary File 8.** *op* and *cp* concatenated coding sequences of the 32 genes located in the candidate region (fasta format).

*See the attached Excel file SupplementaryFile8.txt.*

**Supplementary File 9.** *Acyrthosiphon pisum* populations collected on *Medicago sativa*. Sampling information and observed sequencing depth are provided.

| Reproductive mode | Location | Latitude | Longitude | Pop ID | Number of lineages per pool | Libraries ID | Mean depth per library |
| --- | --- | --- | --- | --- | --- | --- | --- |
| Cyclical parthenogenetic (CP) populations | Saint-Prex (Switzerland) | 46°28’ N | 6°26’ E | Sl | 21 | Sl02  Sl08 | 20.4  16.2 |
|  | Ranspach (France) | 48°01’ N | 7°33’ E | Vl | 20 | Vl03  Vl09 | 18.8  16.2 |
|  | Mirecourt (France) | 48°16’ N | 6°06’ E | Mil | 20 | Mil01  Mil07 | 17.3  17.9 |
| Obligate parthenogenetic (OP) populations | Castelnaudary (France) | 43°19’ N | 1°57’ E | Cast | 14 | Cast04  Cast10 | 18.8  15.6 |
|  | Gers  (France) | 43°57’ N | 0°22’ E | Gers | 14 | Gers05  Gers11 | 15.8  15.1 |
|  | Lusignan  (France) | 46°24’ N | 0°04’ E | Lus | 14 | Lus06  Lus12 | 15.8  17.2 |

**Supplementary File 10.** Principal component analysis score plot of the first two components calculated on allele frequencies for the 12 libraries (six populations, with two replicates each). OP populations are shown in blue and CP populations in red. The first axis (that accounts for 29% of the variance) separates populations by reproductive mode and the second axis discriminates mainly the different OP populations (14% of variance explained).

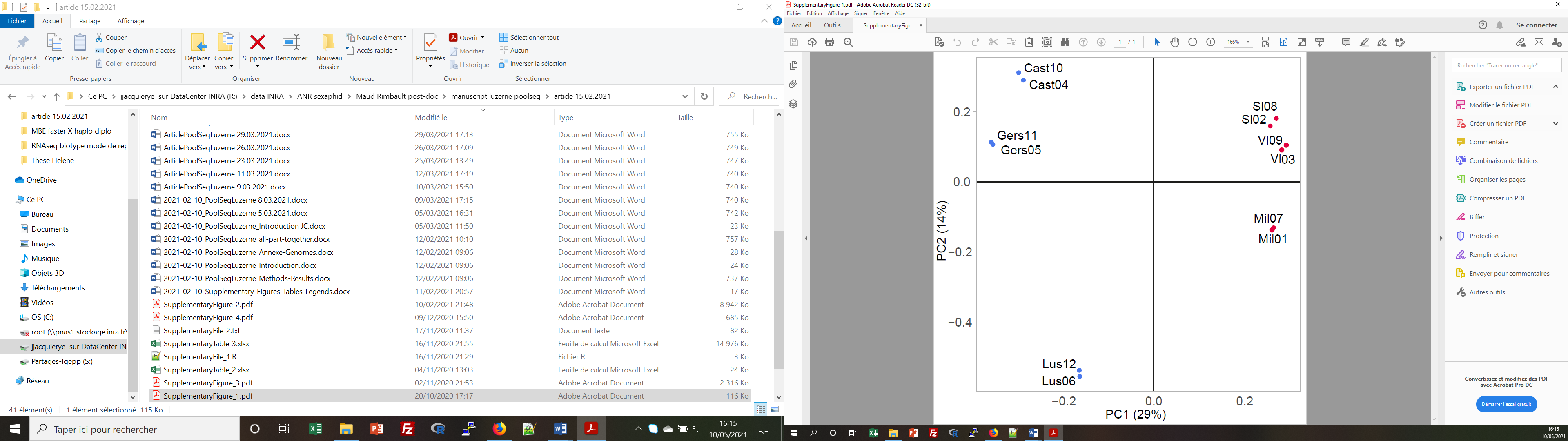
